# A CRISPR screen of HIV dependency factors reveals *CCNT1* is non-essential in T cells but required for HIV-1 reactivation from latency

**DOI:** 10.1101/2023.07.28.551016

**Authors:** Terry L Hafer, Abby Felton, Yennifer Delgado, Harini Srinivasan, Michael Emerman

**Affiliations:** Molecular and Cellular Biology Graduate Program, University of Washington, Seattle, WA 98195, USA; Divisions of Human Biology and Basic Sciences, Fred Hutchinson Cancer Center, Seattle, WA 98109, USA; Bioinformatics Shared Resource, Fred Hutchinson Cancer Center, Seattle, WA 98109, USA

## Abstract

We sought to explore the hypothesis that host factors required for HIV-1 replication also play a role in latency reversal. Using a CRISPR gene library of putative HIV dependency factors, we performed a screen to identify genes required for latency reactivation. We identified several HIV-1 dependency factors that play a key role in HIV-1 latency reactivation including *ELL*, *UBE2M*, *TBL1XR1*, *HDAC3*, *AMBRA1*, and *ALYREF*. Knockout of Cyclin T1 (*CCNT1*), a component of the P-TEFb complex important for transcription elongation, was the top hit in the screen and had the largest effect on HIV latency reversal with a wide variety of latency reversal agents. Moreover, *CCNT1* knockout prevents latency reactivation in a primary CD4+ T cell model of HIV latency without affecting activation of these cells. RNA sequencing data showed that CCNT1 regulates HIV-1 proviral genes to a larger extent than any other host gene and had no significant effects on RNA transcripts in primary T cells after activation. We conclude that CCNT1 function is redundant in T cells but is absolutely required for HIV latency reversal.

## Introduction

The existence of an activatable latent reservoir is a key barrier to virus elimination in people living with HIV as cells which harbor an integrated latent proviral genome persist in the presence of antiretroviral treatment. The multifaceted nature of HIV latency suggests a combination of methods and approaches will need to be used to effectively reduce this reservoir. Factors that ultimately block HIV-1 transcription including host epigenetic silencing mechanisms, blocks to transcription initiation and transcription elongation all contribute to a silent, or nearly silent, HIV reservoir.

The “shock and kill” approach to reservoir reduction involves using latency reversal agents (LRAs) to promote viral transcription and viral reactivation in the latent reservoir and then eliminating those reactivated cells using immunological approaches or methods that rely on recognition of newly synthesized viral proteins (1–3). The shock and kill approach is attractive in that it seeks to eliminate the latent reservoir by killing cells harboring transcriptionally-competent proviral sequences. However, these LRAs must target a broad range of proviruses with highly-variable epigenetic and gene expression contexts in different cells and tissues (4, 5). Another strategy, called “block and lock”, involves targeting factors that are required for HIV replication in order to prevent viral reactivation (6, 7). Such approaches rely on molecules called Latency Promoting Agents (LPAs) that seek to lock the HIV promoter into a permanently silenced state. For instance, didehydro-Cortistatin A (dCA) inhibits Tat/TAR interaction and therefore enforces latency by inhibiting Tat transactivation (8). Other approaches have used siRNAs to target the LTR and prevent transcription of proviral genes which can lead to epigenetic silencing on recruitment of histone modifying complexes to the LTR region (9, 10). Thus far, only one block-and-lock drug, ruxolitinib – a JAK/STAT inhibitor, has made it to a clinical Phase 2a study (11). Both “shock and kill” and “block and lock” therapeutic approaches will likely involve manipulation of multiple arms of HIV latency for a desired outcome, and therefore a more comprehensive understanding of these mechanisms is an important consideration for approaches to eliminate the latent reservoir and achieve a functional HIV cure.

We previously performed a CRISPR screen using a novel system called Latency HIV-CRISPR to identify host genes involved in epigenetic control that maintain latency (12). In this screen, knockout of genes promotes reactivation from latency, suggesting that these host genes normally function to repress HIV-1 transcriptional activation. In the present study, we modified this system to identify host genes that are required for HIV-1 to reactivate from latency, i.e. are necessary for HIV-1 to come out latency. We hypothesized that a subset of host genes that HIV requires for replication, called HIV dependency factors, would also be required for reactivation from latency. Our goal was to identify proteins whose function is more important for HIV-1 reactivation than for normal T cell biology.

Transcription of HIV-1 is dependent on several host mechanisms, with the P-TEFb complex being a key component that interacts with a viral protein, Tat, and a viral RNA element, TAR, to allow for transcription elongation. Both HIV-1 and host genes use CCNT1 and CDK9 in the P-TEFb complex in order to enable transcription elongation (13). CCNT1 has a paralog – CCNT2 – which also forms the P-TEFb complex (14) and *in vitro* studies have shown that another host protein CCNK, can also interact with CDK9 to form the P-TEFb complex (15). However, while HIV-1 Tat viral protein binding sites are conserved in CCNT1 and CCNT2 only the CCNT1-Tat complex can bind with the viral TAR RNA in order to recruit P-TEFb to the LTR (16).

Here, we performed a CRISPR-Cas9 screen using the Latency HIV-CRISPR technique (12) for factors necessary for HIV-1 to be released from latency in the presence of a combination of LRAs. We used a custom CRISPR guide library, called the HIV dependency factor gene library (HIV-Dep), that had been previously used to identify novel host dependency factors across multiple HIV strains(17). We identified and validated factors important in latency reactivation including *ELL1*, *TBL1XR1*, *UBE2M*, *HDAC3*, *AMBRA1*, and ALYREF. Cyclin T1 (*CCNT1*), which forms the P-TEFb transcriptional elongation complex with Cyclin-dependent Kinase 9 (*CDK9*) was the top gene hit in two J-Lat models in our screen. We found that Cyclin T1 is essential for reactivation from latency in J-Lat cells as well as in a primary T cell model of HIV latency using a broad range of LRAs. *CCNT1* knockout had no effect on cell proliferation in the J-Lat model, and did not affect activation through the T cell receptor in primary CD4+ T cells. Moreover, we performed bulk RNA sequencing on *CCNT1* knockouts and found HIV-1 genes were the most depleted relative to wild-type *CCNT1* over any host gene in J-Lat cells, whether or not treated with an LRA. RNA sequencing in uninfected primary T cells knocked out for *CCNT1* showed very few changes in host cell transcript expression. Together, our findings show that some HIV-1 dependency factors are more important for HIV replication and reactivation than for host cell biology and suggest that *CCNT1* could be a promising therapeutic target for silencing HIV-1 into deeper latency. To that end, other genes uncovered in our screen may also be worth exploring further as factors for a block and lock mechanism for HIV.

## Results

### A Latency HIV-CRISPR Screen of HIV Dependency Factors to Identify Latency Reversal Factors

We recently developed and validated a CRISPR sublibrary of guide RNAs targeting host genes important for HIV replication across multiple strains (the HIV dependency factor or HIV-Dep library). The HIV-Dep library has guides targeting 525 genes represented by 8 guides targeting each gene and 210 non-targeting controls (NTCs) (17). A MetaScape analysis (18) of the HIV-Dep library shows the most enriched gene ontology is chromatin organization, followed by several processes involving gene expression, DNA metabolism, and viral infection pathways (Figure 1A). Genes in many of these categories were previously validated to be important in acute HIV-1 infections (17). We hypothesized that a subset of these HIV dependency factors are also necessary for activation of HIV from latency. Thus, to investigate host genes that are required for reversal of HIV-1 latency, we performed a CRIPSR screen using a modification of the HIV-CRISPR system (12, 19, 20) (Figure 1B). Briefly, this screen in the context of latency reversal relies on transducing latently infected Jurkat T cells (J-Lats) with an HIV-CRISPR lentiviral vector containing a library of sgRNAs. The sgRNAs are flanked by a Ψ-packaging signal, allowing the guides to be packaged into budding virions. We employed this modified latency HIV-CRISPR assay to identify factors important for latency reactivation using two different J-Lat models that contain independently-derived integration sites; J-Lat 10.6 and J-Lat 5A8. The goal for this screen was to treat the cells with activating doses of LRAs, deep sequence the supernatant containing the guides compared with the gDNA knockout pool. In contrast to a previous HIV-CRISPR screen where we examined epigenetic factors whose knockout would activate HIV from latency by analyzing guides enriched in the viral supernatant (Figure 1B, scenario 1) (12), in the present screen the expectation is that genes required for reactivation from latency would be depleted in the viral supernatant relative to the genomic knockout pool (Figure 1B, scenario 2).

**Figure 1.**
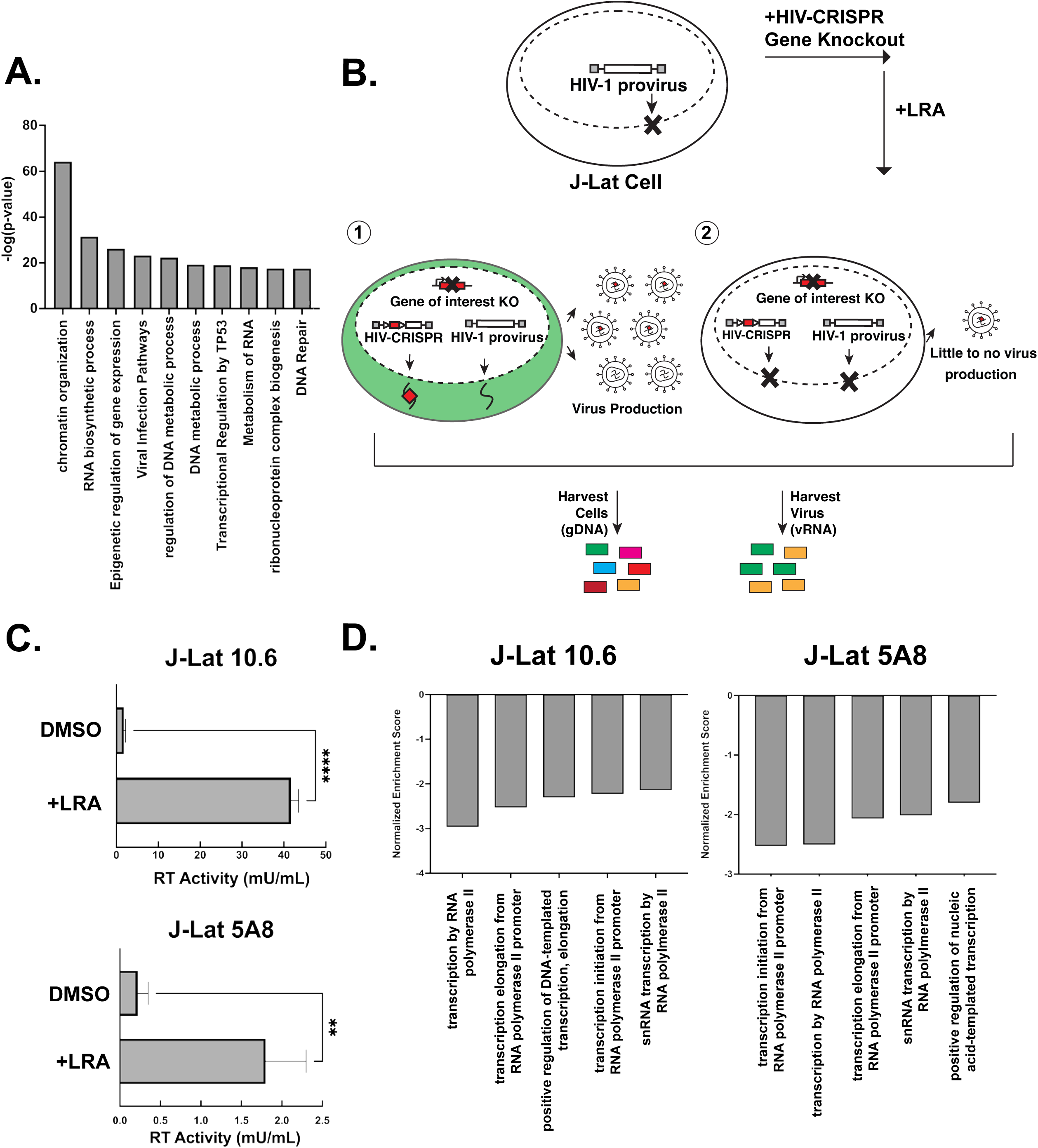
A Latency HIV-CRISPR Screen to identify factors required for latency reversal. **(A)** A Metascape analysis of the genes in the HIV-Dep gene library is shown, with enriched pathways on the x-axis and statistical significance on the y-axis. **(B)** Overview of latency HIV-CRISPR screen of HIV Dependency Factors. The HIV-CRISPR vector has intact 5’ and 3’ LTRs and can be packaged by HIV-1 after integration (19) J-Lat cells were transduced with an HIV-CRISPR library of genes of HIV-1 dependency factors, selected for integration by puromycin selection, and treated with a latency reversal agent (LRA). Viral RNA (vRNA) and genomic DNA (gDNA) are harvested at the end of the experiment. Guides corresponding with genes that do not affect reactivation from latency are packaged in virions and enriched in the supernatant relative to the genomic DNA pool (scenario 1, left). For genes that are important for latency reactivation after treatment of cells with an LRA, these guides will be depleted in the viral supernatant relative to the genomic DNA knockout library (scenario 2, right). **(C)** Supernatant from J-Lat cells transduced with the HIV-DEP gene library were measured for Reverse Transcriptase (RT) activity after treatment with the LRA combination AZD5582 (1 nM) and I-BET151 (2.5 uM). Error bars represent technical triplicates, unpaired t-test was used for statistical analysis. p-value < 0.01 = **, < 0.0001 = **** **(D)** MAGEcKFlute (22) was used to analyze screen results of the depleted genes. The normalized enrichment score is on the y-axis (negative because guides to these genes are depleted from the viral supernatant) and the x-axis is the biological processes.

J-Lat cells transduced with the HIV-Dep library were treated with low doses of the non-canonical NF-κB inhibitor AZD5582 (1 nM) and the pan-bromodomain inhibitor I-BET151 (2.5 uM), which led to significant increases in viral production as measured by reverse transcriptase activity (Figure 1C). Previous studies had also shown that this combination of LRAs is synergistic in the J-Lat model of latency reversal (21). After deep sequencing the viral supernatant and genomic DNA pool, we used MAGEcK analysis in order to compare the guides enriched or depleted in the supernatant with the genomic knockout pool to identify those genes depleted in the supernatant (Supplemental File 1). We generated a gene set enrichment analysis (22) of our most depleted hits and found the top five enriched pathways in both J-Lat 10.6 and J-Lat 5A8 were related to transcription (Figure 1D). Furthermore, we also saw pathways for RNA splicing and polyadenylation. This is consistent with transcriptional regulation being one of the major axes of host control that underly release of HIV-1 from latency. We conclude that our screen can identify and enrich for gene pathways that are relevant for release of the HIV-1 provirus from latency in the presence of AZD5582 and I-BET151 combination treatment.

To understand the role that HIV dependency factors play in terms of latency reactivation, we compared our screens with previous HIV-CRISPR screens that were aimed at identifying factors required for HIV replication in Jurkat cells (17). A z-score analysis was used as a measure of how depleted genes were in each of the screens and to allow for a cross-comparison regardless of the magnitude of depletion of each guide. Sorting the mean z-score for HIV-1 replication (marked as LAI in Figure 2A) shows that the most depleted genes are *CXCR4* and *CD4* which are essential for HIV replication but not for latency reactivation (Figure 2A, left). This is expected since J-Lat cells are already infected with HIV-1. Other factors that scored highly in the HIV-1 replication screen, but not in the present HIV latency screen include genes of unknown function in the HIV lifecycle such as *ATP2A2* and *SS18L2* (Figure 2A, left). In contrast, nearly all of the most depleted factors in the HIV latency screens were also highly depleted in the HIV replication screen (Figure 2A, right, sorted by most depleted in the HIV latency screens; see Supplemental 1 for the complete list of Z-scores). We conclude that a subset of HIV dependency factors are required for reactivation from latency.

**Figure 2.**
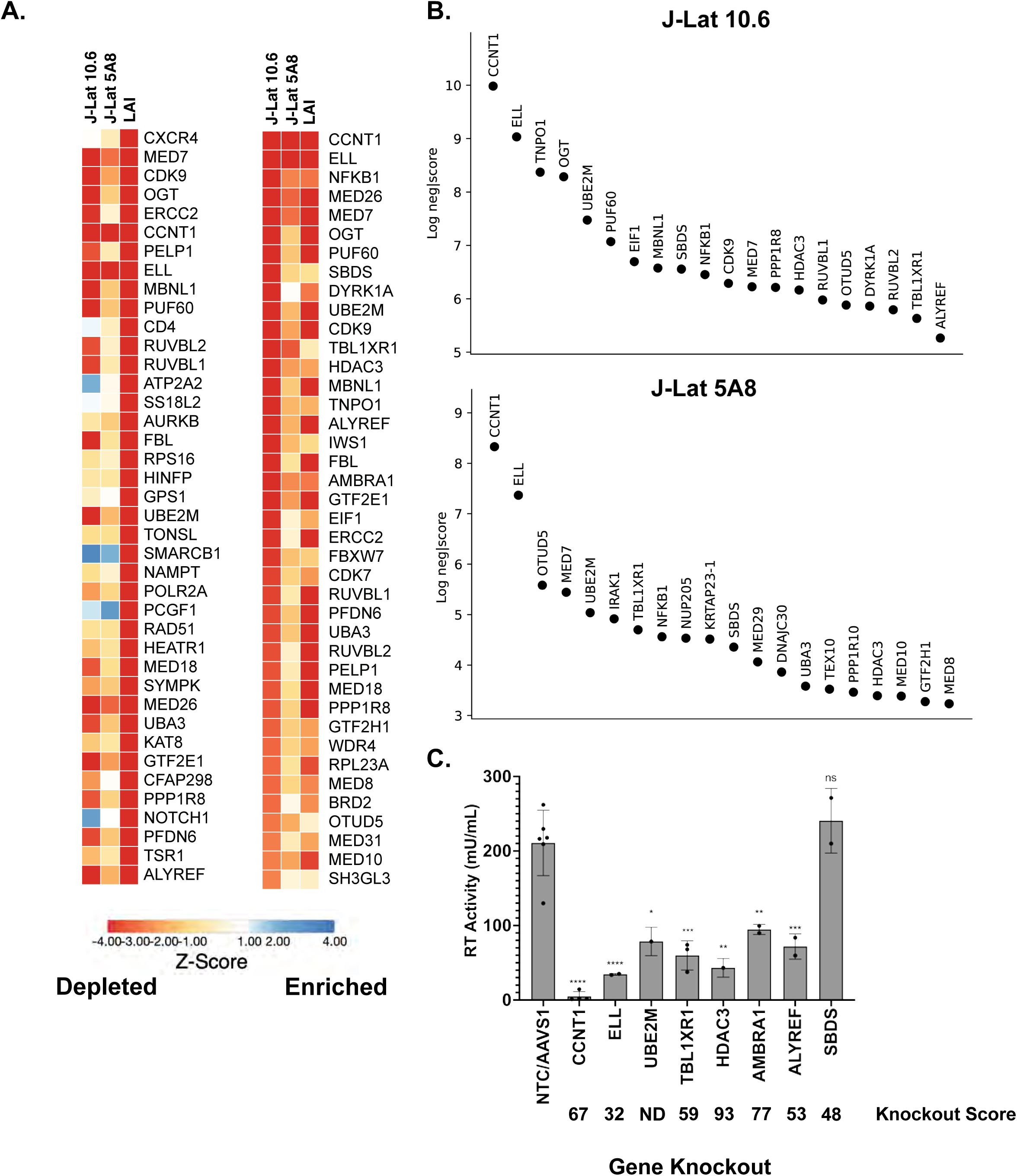
Analysis and Validation of Top Hits from HIV-CRISPR screen. **(A).** Z-score analysis of the depleted versus enriched guides across multiple screens. J-Lat 10.6 and J-Lat 5A8 are screens from this study, whereas LAI represents Jurkat cells infected with an LAI strain of HIV-1 from previous screen performed using the same gene library in Jurkat cells to identify HIV Dependency Factors (17). Z-scores are sorted by the most depleted genes in the LAI screen (left panel) and by the most depleted genes in the J-Lat 10.6 line from this study (right panel). The mean z-score of two replicates each of J-Lat 10.6 and J-Lat 5A8, and of four replicates of the LAI screen is shown. Most depleted genes are red and most enriched genes are blue. Z-scores were that were less than -4 were capped at -4 in the heat map. **(B)**. The top 20 most depleted hits from each J-Lat line in ranked order are shown. **(C)** Selected hits from the screen were tested by performing gene knockouts (x-axis), treating with the LRA combination AZD5582/I-BET151, and assayed for reverse transcriptase activity. Gene knockouts were performed using a lentiviral knockout approach and/or an electroporation with Cas9 and RNPs. Each point represents a single lentiviral or electroporation knockout experiment done in triplicate. An average of RT activity from two guides targeting each gene was taken for lentiviral knockouts, and the electroporation knockouts included three individual guides targeting each gene. The ICE gene knockout score for each experiment was averaged and is shown below each gene on the x-axis. Statistical analysis was performed using a two-way ANOVA and Šídák’s multiple comparisons test to measure the difference in latency reactivation between each gene knockout relative to NTC/AAVS1 control. p-value ≥ 0.05 = ns (not significant), < 0.05 = *, < 0.01 = **, < 0.001 = ***, < 0.0001 = ****. NTC/*AAVS1* controls are combined; each dot represents either an *AAVS1* or NTC control for an individual experiment. Each experiment (dot) has 3 technical replicates: NTC/AAVS1, n=6 experiments, 3 replicates each; *CCNT1*, n = 4 experiments, 3 replicates each; *ELL*, n = 2 experiments, 3 replicates each; *UBE2M,* n = 1 experiment, 3 replicates each; TBL1XR1, n = 3 experiments, 3 replicates each; *HDAC3*, 1 experiment, 3 replicates each; *AMBRA1* n = 2 experiments, 3 replicates each; *ALYREF*, 2 experiments, 3 replicates each; *SBDS* n = 2 experiments, 3 replicates each.

We chose to validate a subset of the hits in the HIV latency screen that were among the top twenty ranking hits and were shared hits in both J-Lat 10.6 and J-Lat 5A8 cells (Figure 2B, complete list of the screen in Supplemental File 1) by electroporating Cas9 ribonucleoprotein complex (RNP) complex containing 3 unique guides against each gene or by lentiviral transduction of single guide RNAs. We tested *CCNT1*, *ELL*, *UBE2M, TBL1XR1, HDAC3*, *AMBRA1, ALYREF, and SBDS* (Figure 2C). As a negative control we included guides targeting the adeno-associated virus integration site 1 (*AAVS1*) “safe harbor” locus, a gene whose disruption does not adversely affect the cell (23), or a non-targeting control (NTC). Knockouts were validated by genomic sequencing. In the J-Lat 10.6 line we found that there is reduced reactivation in *CCNT1*, *ELL*, *UBE2M, TBL1XR1, HDAC3*, *AMBRA1,* and *ALYREF* knockouts relative to non-targeting controls and guides targeting a safe harbor locus, *AAVS1* (Figure 2C). We did not see a significant effect in the *SBDS* knockout cells, but interestingly *AMBRA1* and *ALYREF* which were less depleted than *SBDS* in the J-Lat screens did show a phenotype. However, the strongest effect on preventing HIV latency reversal was the knockout of *CCNT1* which was also the top hit in our screen. We conclude that the screen is able to identify genes that are key for latency reactivation in the J-Lat models.

### Cyclin T1 is essential for Latency in both J-Lat and primary T cells

Cyclin T1 (CCNT1) is a well characterized regulator of HIV transcription that binds to the viral protein Tat and TAR (24–26) and was the top hit for both J-Lat models. Additionally, CDK9 which binds to Cyclin T1 in order to form the positive transcription elongation factor complex (P-TEFb) is substantially depleted in both cell lines. In order to explore this hit further across a broader range of LRAs, we generated clonal knockout lines of *CCNT1* in the J-Lat 10.6 cell line. The clonal knockouts are completely abrogated of CCNT1 expression as shown by Western blotting and by sequencing of genomic DNA (Figure 3A, left). Moreover, we did not see an upregulation of CCNT2, a paralog of CCNT1 that also binds CDK9 as part of the host P-TEFb complex (14, 16) (Figure 3A, right).

**Figure 3.**
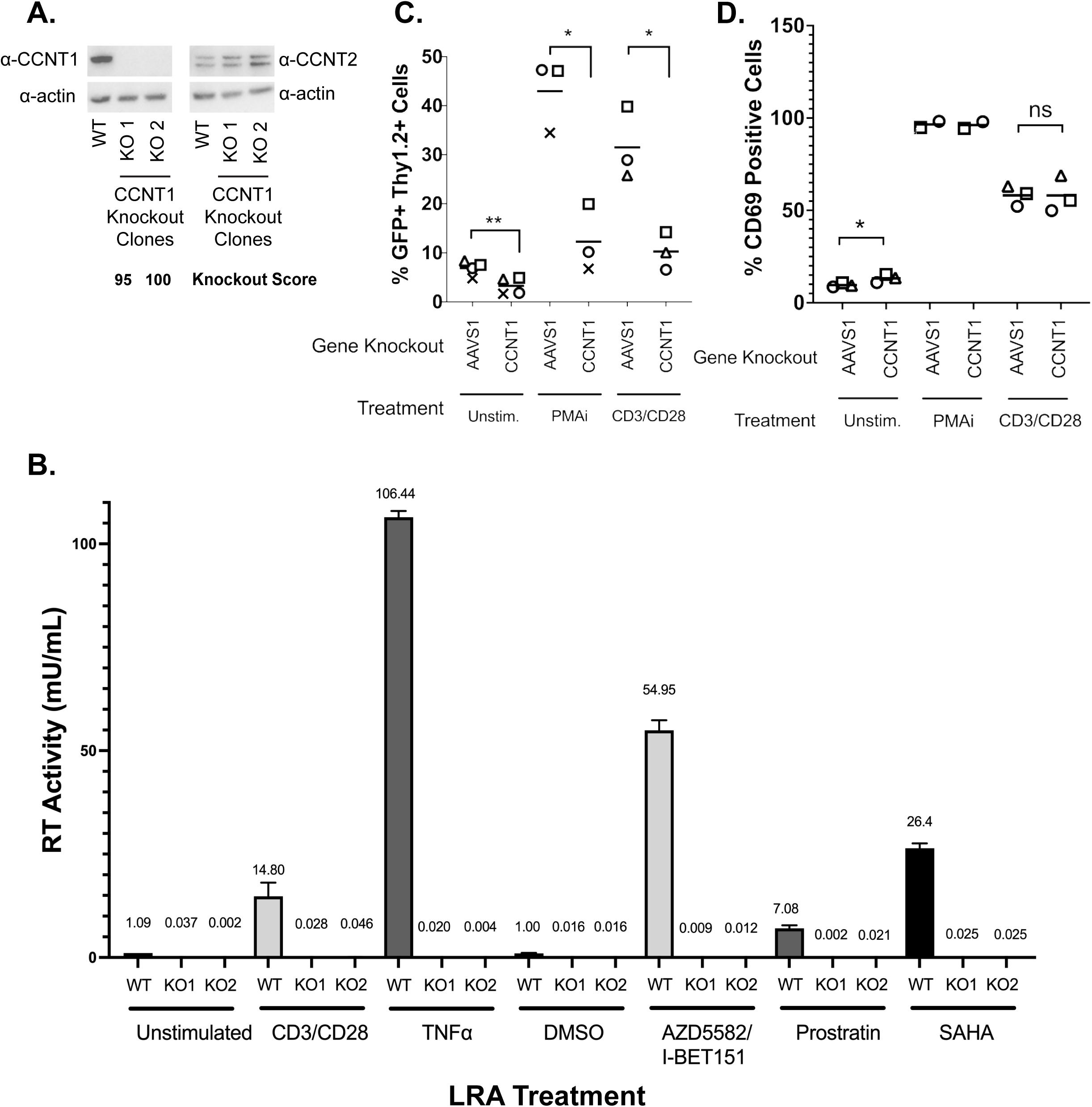
*CCNT1* is required for reactivation of HIV-1 from latency in Jurkat T cells and primary CD4+ T cells from healthy donors. **(A)**. Western blot of cell lysates of J-Lat 10.6 either wild-type or clonally knocked out for CCNT1 is shown, with two separate knockout clones. Actin was used as loading control. Left: CCNT1 antibody is shown, Right: CCNT2 antibody is shown. ICE Knockout scores are shown for each knockout clone of *CCNT1* **(B).** J-Lat 10.6 cells wild-type for CCNT1 and the two clones knocked out for *CCNT1* were treated with the LRAs shown on the bottom. The mean of RT activity in the supernatant 24 hrs after LRA treatment is shown on the Y axis and above each bar. Averages and standard deviation of the experiment done in triplicate is represented **(C).** Primary CD4+ T cells from three different healthy donors were infected with a dual-reporter virus that monitors cells active and latent infection (Thy 1.2, CD90 marker) and actively transcribing provirus (GFP marker). Cells were either knocked out for *AAVS1* control or *CCNT1* and either untreated, stimulated with PMAi, or stimulated with anti-CD3/anti-CD28 antibodies at the end of latency establishment. Each shape represents an individual donor. **(D)** CD69 expression was monitored with the different LRA treatments. *CCNT1* ICE knockout scores were: 80, 76, 53, and 37 for each of four donors for CD3/CD28 and two donors for PMAi. A paired t-test was used for comparison of *AAVS1* knockout vs *CCNT1* knockout between donors. p-value ≥ 0.05 = ns, < 0.05 = *, < 0.01 = **

HIV latency is a result of a combination of blocks that prevent transcription initiation and elongation, and LRAs target a broad range of these different facets of proviral gene expression. We explored a range of LRAs in the *CCNT1* clonal knockout lines. We found that CCNT1 is necessary for latency reversal with both CD3/CD28 activation and with Tumor Necrosis Factor Alpha (TNFα) cytokine. Reactivating with CD3/CD28 and TNFα are mechanisms that result in the upregulation of NF-κB signaling, a facet that emphasized the transcription initiation component of latency. We therefore explored additional means of reactivation including AZD5582 and I-BET151 together, Prostratin – an activator of PKC and known inducer of P-TEFb activity (27, 28) – and SAHA/Vorinostat (29), the histone deacetylase inhibitor (HDACi) (Figure 3B). In all treatments, cells wild-type for CCNT1 were able to reactivate, but *CCNT1* knockout prevented latency reactivation with each LRA. We conclude that CCNT1 is essential for reactivation from latency for multiple diverse mechanisms of latency reversal in J-Lat cells.

We also investigated the role of CCNT1 in latency reactivation in primary CD4+ T cell lymphocytes isolated from healthy donors. We first activated and infected peripheral blood CD4+ T cell lymphocytes with an HIV-1 dual-reporter virus previously described (12); the first marker is a destabilized GFP reporter is a marker of active provirus expression. The destabilized GFP has a short half-life and thus is indicative of active expression of the provirus. The second marker, Thy1.2 (mouse CD90) viral reporter is a cell surface marker that allows for us determine cells that have, at one point, been infected. This cell surface marker has a slow turnover and persists over the latency establishment period, and thus marks cells that have been infected with the dual-reporter virus, but may not be actively producing virus. After infection with dual reporter virus, infected cells were knocked out by electroporation with Cas9 and gRNA for *CCNT1* or control *AAVS1*. Cells were cultured for an additional two weeks to enter latency, and then measured for the capability for latency reactivation after LRA treatment as determined by flow cytometry for dual positive GFP and CD90 expression (Figure 3C and Figure S1).

We tested knockouts from three independent donors with the potent LRA combination phorbol 12-myristate 13-acetate (PMA) and ionomycin as well as with CD3/CD28 antibody costimulation (Figure 3C for all donors, Supplemental Figure 1 for the gating of one donor as an example). In control *AAVS1* knockout we found that there is an increase in the percentage of total cells that are both Thy1.2+ and GFP+ on treatment with PMAi or CD3/CD28 co-stimulation indicating an increase in cells that have active transcription of viral genes (5.46% without LRA, 39.7% with LRA) (Figure 3C). In contrast, the *CCNT1* knockouts had a stark reduction in Thy1.2+ and GFP+ cells on treatment with PMAi and CD3/CD28 co-stimulation relative to *AAVS1* knockout (Figure 3C, S1). We also noted that there is a modest reduction of Thy 1.2+ GFP+ cells in the *CCNT1* knockout that have not been treated with PMAi or CD3/CD28 costimulation. This is consistent with our previous result in clonal knockouts in J-Lat cells suggesting that minimal levels of HIV-1 transcription that occur in latent cell populations are lower in *CCNT1* knockouts. We conclude that Cyclin T1 is an essential gene for latency reactivation.

To exclude the possibility that Cyclin T1 blocks the ability for CD4+ T cells to activate, as well as ensure T cell activation is occurring properly in our experiments, we simultaneously stained cells for the early activation marker CD69. PMAi and CD3/CD28 co-stimulation both show a significant degree of activation over unstimulated cells. We saw no significant change between *AAVS1* and *CCNT1* knockout in any of the conditions (Figure 3D). We conclude that *CCNT1* is key for latency reactivation in primary CD+4 T cells but does not affect the ability of these cells to be activated upon stimulation.

### Cyclin T1 is non-essential in T cells and regulates host genes to a much lesser extent than it regulates HIV-1

Given that P-TEFb has been reported to be required for transcription elongation of many host genes (30), we were initially surprised that knockout of *CCNT1* is viable. However, we did not see a drastic change in cell growth measured over a span of nine days (Figure 4A). This led us to broadly investigate the role of Cyclin T1 in transcription in T cells by performing bulk RNA sequencing of J-Lat 10.6 cells and two independent clonal knockouts of *CCNT1* in the J-Lat 10.6 cells either without an LRA, or treated with TNFα. As a control, we first compared the RNA sequencing data from wild-type J-Lat 10.6 line that has been treated with TNFα, versus the J-Lat 10.6 line (*CCNT1* is wild-type in both cases). HIV-1 transcripts are among the most significantly upregulated genes in the TNFα treatment for wild-type (Figure 4B). We also see upregulation of *PGLYRP4, RELB, and BCL3*, which are genes related to NF-κB signaling or otherwise known to be upregulated by TNFα (Figure 4B) (31–33). We next examined how HIV-1 and host gene transcripts are affected in TNFα treated cells that have *CCNT1* knocked out relative TNFα treated J-Lat 10.6 cells that are wild-type for *CCNT1* (Figure 4C). Strikingly, RNA transcripts related to HIV-1 genes in *CCNT1* knockout are the most depleted transcripts over any host gene, relative to wild-type *CCNT1* (Log_2_(FC) = -10.92) (Figure 4C). Even in the absence of LRA, we find that HIV-1 transcripts are the most depleted relative to other host genes (Log_2_(FC) = -9.29) when comparing *CCNT1* knockout versus wild-type (Figure 4D). Thus, basal transcription of HIV-1 transcripts that occur in J-Lat lines are highly dependent on Cyclin T1. Regardless of TNFα treatment, the host genes that were highly depleted in *CCNT1* knockout included *FAM222A-AS*, *GGTLC1*, *MYO10, NETO1*, and *ZBTB16*. Notably, we did not find significant upregulation of *CCNT2* transcripts in the CCNT1 knockout versus wild-type (Log_2_(FC) = 0.078) or in the LRA treated cells (Log_2_(FC) = 0.139). Nonetheless, *CCNT1* knockout affects the HIV-1 provirus far more than any other transcriptional unit in the J-Lat cells.

**Figure 4.**
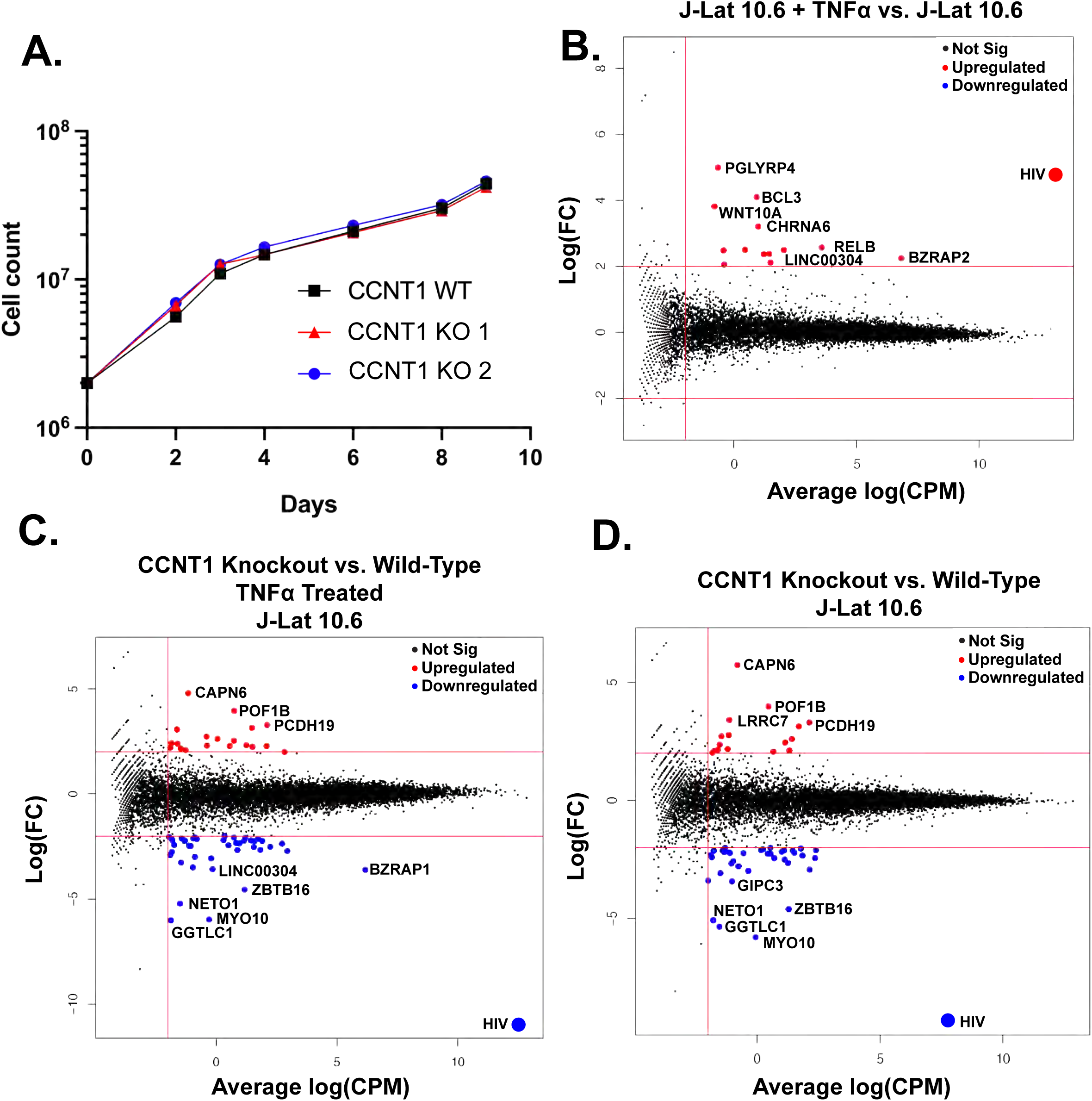
Cell proliferation and RNA sequencing analysis of *CCNT1* knockouts in J-Lat 10.6 cells. **(A).** Cell counts were monitored over a span of nine days in J-Lat 10.6 cells in WT or clonally knocked out *CCNT1* cells. The average of three experimental replicates are shown with standard deviation. **(B-D).** Log_2_ FC (fold-change) is plotted on y-axis with the average Log_2_ CPM (counts per million) across technical replicates on the x-axis. Red lines on the signify genes that have an average Log_2_ CPM > -1, and a |Log_2_ FC | > 2. Red dots signify upregulated genes whereas blue genes signify downregulated genes for each comparison. **B)** Differential gene expression of J-Lat 10.6 with TNFα treatment versus J-Lat 10.6 (untreated) is shown. **C).** J-Lat 10.6 *CCNT1* KO cells (two independent clones each tested in technical triplicate and averaged) versus the J-Lat 10.6 wild-type cells – both were treated with the LRA TNFα and gene expression comparison is shown. **D).** J-Lat 10.6 *CCNT1* KO cells versus wild-type *CCNT1* differential gene expression is shown – neither cell line was treated with an LRA.

We further investigated the effect of *CCNT1* knockout on uninfected primary CD4+ T cells. *CCNT1* was knocked out by electroporation of *CCNT1* guides complexed with Cas9 in three independent donors and the knockout was validated to be over 90% by sequence analysis (Supplemental File S2). The *AAVS1* locus was knocked out in parallel as a control. Similar to the primary cell latency model (Figure 3C), we found that the *CCNT1* knockout did not affect expression of the CD69 activation marker after treatment with anti-CD3/anti-CD28 beads (Figure 5A). As expected, comparison of RNA sequencing on primary cells stimulated with anti-CD3 and anti-CD28 antibodies versus unstimulated cells shows dramatic upregulation and downregulation of genes (Figure 5B); for example, there is upregulation of IL31 which is a cytokine known to be upregulated by activated T cells (34). However, the same RNA-seq analysis of *AAVS1* knockout cells compared to *CCNT1* knockout cells upon stimulation with anti-CD3/anti-CD28 beads shows that *CCNT1* knockout cells have the same expression profile as the control knockout cells, i.e. there are no significant differences in upregulated or downregulated genes in the comparison (Figure 5C) when *CCNT1* is knocked out. We also compared RNA expression profiles of the *CCNT1* knockout cells with the controls *AAVS1* knockout cells in the absence of anti-CD3 and anti-CD28 stimulation, and again find very few genes which are upregulated or downregulated (Figure 5D). In addition, the magnitude of these gene expression changes was minimal. As an example, the most enriched gene for *CCNT1* knockout compared to *AAVS1* knockout has a -log_2_FC less than 2, and the most depleted gene has a -log_2_FC greater than -2 (Figure 5D). Thus, we conclude that there are minimal changes in gene expression when *CCNT1* is knocked out in primary CD4+ T cells with and without T cell receptor stimulation. Together, we conclude that *CCNT1* does not play an essential role in peripheral primary CD4+ T cells.

**Figure 5.**
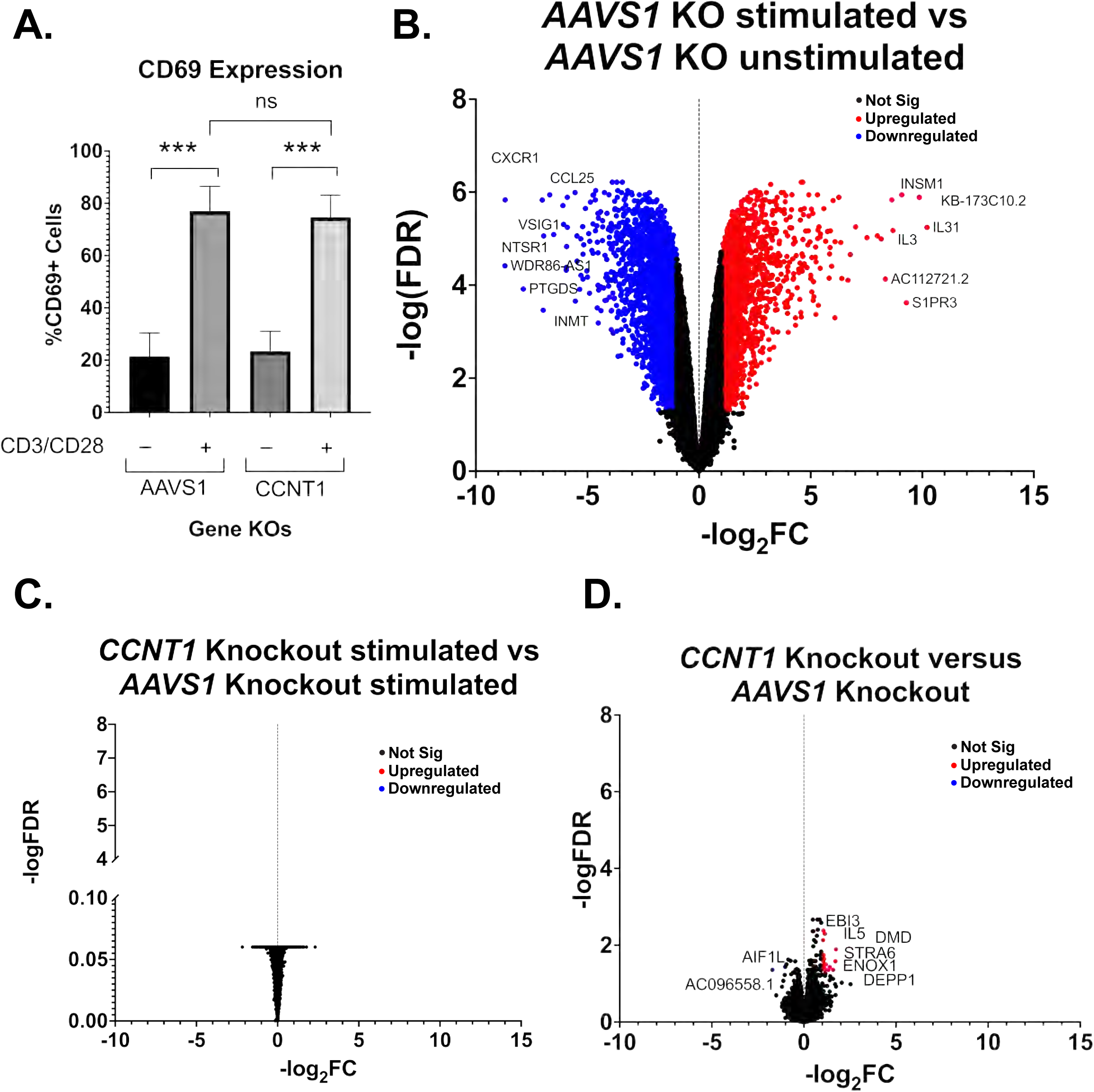
Primary T cells transcripts are largely unaffected by *CCNT1* knockout. **(A)** Uninfected CD4+ T cells from three donors were knocked out for *AAVS1* or CCNT1, and then treated with CD3/CD28 co-stimulation. Cells were analyzed by flow cytometry to measure CD69 expression. On left, one representative donor is shown. On right, a summary of CD69 expression in AAVS1 knockout versus CCNT1 knockout from all three healthy donors is shown. One-way ANOVA was used for analysis with Dunnett’s multiple comparison tests. P (**B-D).** Volcano plots of primary CD4+ T cell RNA sequencing data is shown, with -log_2_FC shown on the x-axis and -log(FDR) on the y-axis. RNA was isolated from three biological replicates. A FDR = 0.05 was used as a cutoff for significance, and the cutoff for significant gene expression was |Fold-Change| > 1. A subset of genes for each condition are marked that have significance. **(B)** Differential gene expression between *AAVS1* knockout stimulated with CD3/CD28 versus unstimulated is shown. **(C)** A comparison of *CCNT1* versus *AAVS1* knockout is shown, and both were stimulated with anti-CD3/anti-CD28 antibodies. **(D)** *CCNT1* versus *AAVS1* knockout is shown, and neither of these are stimulated with anti-CD3/anti-CD28 antibodies. p-value ≥ 0.05 =, < 0.001 = ***

## Discussion

We used an HIV-CRISPR screening approach to identify host genes required for activation of HIV from latency starting from the hypothesis that a subset of host genes previously identified as being necessary for HIV replication are also necessary for HIV reactivation from latency. Among the genes identified include many genes involved in transcription elongation, transcription initiation and protein degradation. The top hit in our screens was Cyclin T1 (*CCNT1*) which we show is essential for reactivation from latency across a wide range of latency reversal agents of different mechanisms of action, as well as in primary T cells. In contrast, *CCNT1* appears to be redundant with other host genes for normal transcriptional regulation in T cells and is therefore an attractive target for specifically silencing integrated HIV-1 proviruses.

### Cyclin T1 is much more important for HIV latency reversal than for T cell biology *in vitro*

Despite the described role of Cyclin T1 and the P-TEFb complex in host gene transcription, we were able to generate knockout clones of *CCNT1* without affecting cell growth and viability. We also did not see a significant upregulation of CCNT2 protein expression. Collectively, we interpret our results to mean that *CCNT1* is dispensable in T cells and that *CCNT2* or *CCNK* may compensate for the loss of *CCNT1*. One model is that there are redundant mechanisms that govern transcription elongation of host genes. Previous work on *CCNT1* and *CCNT2* knockouts in mice illustrated unique phenotypes, initially suggesting the possibility that these two genes have separate functions despite both being able to form the P-TEFb complex (35, 36). RNA sequencing of CCNT1 and CCNT2 knockdowns by another group using shRNA in HeLa cells also suggested these two proteins are regulating different sets of genes (37). However, while CCNT1 had very large effects on HIV-1 transcripts, we found that CCNT1 has minimal effects on host gene transcription in Jurkat cells. We did observe a modest downregulation of several host genes including *GGTLC1*, *MYO10*, *NETO1*, *ZBTB16*, and *BZRAP1*. GGTLC1 is a metabolic enzyme and member of the gamma-glutamyl transpeptidase family, of which there are several paralogs (38). Myo10 is an unconventional myosin that associates with actin and filopodia. This gene has ubiquitous but low expression across tissues (39), but has been reported to promote HIV-1 infection in human monocyte derived macrophages (40). *ZBTB16* (also known as *PLZF*) is a transcription factor and is known to be important for natural killer T cells, but repressed in non-innate T cells and not upregulated in T cell activation (41). Collectively, we see slight changes in gene expression in J-Lat cells on *CCNT1* knockout that lead to drastic changes in HIV-1 gene expression, but few host genes seem to be affected on knockout.

On the other hand, there were no significant changes in gene expression of *CCNT1* knockout versus *AAVS1* knockout in primary CD4+ T cells activated with CD3/CD28 costimulation. Knockouts of *CCNT1* in primary CD4+ T cells also had little effect on cell viability and cell surface expression of an activation marker, CD69. In the unstimulated condition we see some low magnitude gene expression changes; *AIF1L* is a mildly downregulated gene, and to date there is no clear known function of this gene in T cell biology. In human podocytes, this gene is known to function in actomyosin contractility and thus cells which lack this gene have increased filopodia (42). Upregulated genes include *IL5*, *DMD*, *STRA6*, *ENOX1*, and *DEPP1*. None of these genes are particularly implicated in T cell biology. Mutations in the *DMD* (Dystrophin) gene are implicated in Duchenne’s Muscular Dystrophy, an X-linked recessive disorder. We also saw upregulation of *MYOF* (Myoferlin), a gene whose mutations are associated with muscle weakness (43, 44). An interesting possibility is that *CCNT1* positively and negatively regulates genes associated with muscle function, given we saw an upregulation of these genes implicated in muscle disease, and a downregulation of *MYO10* in the J-Lat 10.6 RNA sequencing data on *CCNT1* knockout.

We reason that while CCNT1 and CCNT2 gene regulation may have tissue-specific contexts; CCNT1 is likely redundant in CD4+ T cells. Data from DepMap indicate that *CCNT1* is classified “strongly selective” indicating there are cell lines in which this gene is more essential, but that *CCNK* is it is considered widely essential in most CRISPR screens (45). We interpret this to mean that the role of *CCNT1* may be redundant in T cells for host gene expression but not for HIV-1 activation. Previous work suggests CCNT1 is targeted by proteasomal degradation in resting CD4+ T cells, and thus CCNT1 protein expression in resting CD4+ T cells is low (46–48), but our data suggests that it is not necessary for T cell activation. While we saw little effect of *CCNT1* knockout on host RNA transcripts in a relevant target cell type for HIV-1 infection, we cannot rule out the possibility that *CCNT1* does play a key role in host biology in more differentiated T cell functions or in other HIV-1 prone cell types including macrophages and glial cells.

### Other hits in the HIV-CRISPR screen

Several genes involved in transcription are among our most depleted genes. Notably, NFKB1 – the transcription factor that binds to 5’ LTR to allow for transcription initiation of proviral genes, is among our top hits. We also see other transcription-related genes depleted in both cell lines. ELL – an elongation factor for RNA polymerase II and component of the super elongation complex – is the second most-depleted hit. We also note that there are several post-translational modifying enzymes that are novel in terms of latency reactivation. The Ubiquitin Conjugating Enzyme E2 M (UBE2M) is highly depleted and is known to be involved in the neddylation pathway, which uses a ubiquitin-like conjugation process. UBA3, which makes up the E1 enzyme of the neddylation conjugation pathway, also is depleted but to a lesser degree. Both of these neddylation genes were also depleted in our previous CRISPR screen on Jurkat T cells to identify dependency factors, and *UBE2M* validated for several strains of HIV (17). Histone Deacetylase 3 (HDAC3) forms a complex with TBL1XR1 as part of the SMRT N-CoR (nuclear coreceptor complex), which regulates modification of histones and gene regulation (49–51). siRNA studies of TBL1XR1 have found redundancy with its paralog TBL1X, whereas HDAC3 was found to be essential. Vorinostat, a commonly used LRA targets HDAC3 along with Class I and Class II HDACs (52). It is unclear why *HDAC3* knockout may prevent latency reactivation, but we reason latency reactivation depends in part on a noncatalytic activity of *HDAC3*.

A genome wide CRISPR screen was previously performed that identified factors important for latency reversal (53). In that study, the authors generated a pool of latently infected cells and performed a whole genome CRISPR knockout screen, treated with a panel of different LRAs, sorted for GFP-cells and identified genes specific for latency reversal as well as common genes required regardless of reactivation approach. In comparing our screens, we find many of our hits are shared with the “common” cluster of genes where they tested TCR cross-linking, TNF-a, PMAi, and AZD5582 as LRAs and identified the common genes required for reactivation: *CCNT1, HDAC3, NFKB1, MBNL1, UBE2M, TBL1XR1, UBA3, AMBRA1, SBDS,* and *MED7*. Thus, despite only screening with AZD5582 and I-BET151, we are able to identify several hits that promote latency reactivation regardless of LRA used. *UBA3* and *UBE2M* are of interest as they are both components of the neddylation pathway (54), and while *NEDD8* is not in our HIV-DEP gene library – the whole genome screen identified *NEDD8* as a hit in their AZD5582 screen (53). In contrast, there are several hits that are depleted and validated in our more targeted screens but not the whole genome screen such as *ELL* and *ALYREF* (Figure 2). Nonetheless, there is overall good agreement between screens, validating the approach of searching for host factors involved in latency through CRISPR screens combined with LRAs.

### HIV Dependency Factors versus host genes necessary for latency reversal

Our initial hypothesis was that HIV-1 dependency factors may play a role in latency reactivation given the importance of transcription in establishing infection and that transcription is a major facet that contributes to latency. Consistent with our hypothesis, we find that a large proportion of genes are important as both HIV dependency factors and as HIV latency reversal factors (Figure 2A). While transcription is the major category of genes in our screens (Figure 1C and D), the factors however span beyond transcription; we find factors involved that are key for reactivation, including *UBA3*, *UBE2M*, *AMBRA1* and *ALYREF*. In contrast, we also observe factors that are important as HIV-1 dependency factors but not in latency reactivation including *ATP2A2*, *SS18L2*, *SMARCB1* and *PCGF1* that were depleted in Jurkat T cells screens but not in J-Lat screens. *ATP2A2* is a Calcium Transporting ATPase that was found to be upregulated during G1/S phase of the cycle by Tat, but its role in the viral life cycle is otherwise unknown (55). Similarly, *SS18L2* was found to be upregulated in HIV-1 in early infection, as found from RNA profiling of CD4+ and CD8+ T cells in people living with HIV-1 versus those who were either nonprogressors or control HIV-1 negative groups (56). SMARCB1 is a component of the SWI/SNF chromatin remodeling complex along with INI1 (Integrase Interactor-1) and is known to play many roles in HIV-1 replication, including integration, transcription and particle maturation (57). *PCGF1* (Polycomb group RING finger protein 1) was also depleted in HIV-1 dependency factor screens, but not in J-Lat screens in this study. Polycomb Group Proteins largely lead to transcriptional repression through methylation of histones, and thus are thought to contribute to HIV-1 latency. This might contribute to the opposite phenotype we see in this study versus infection screens; *PCGF1* may play a role in maintaining latency but is required for establishing infection. An interesting possibility is that PCGF1 is required for infection as it helps to establish a chromatin landscape that leads to either productive transcription at the integrated provirus, or even transcriptional silencing which may ultimately contribute to HIV-1 latency. Collectively, the latency HIV-CRISPR screens can help to narrow down the stage of the viral life cycle dependency factors are playing a role in, but also can give insight into novel latency reversal factors.

### Gene Paralogs in a “Block and Lock” Latency Approach

Our Latency HIV-CRISPR screen in this study revealed our top hit *CCNT1* was able to be knocked out with little effect on cell biology, likely due in part to its paralogs *CCNT2* and *CCNK*. This approach to “block and lock,” whereby a factor is required for viral replication but not for host function, may be a good path forward in further identifying gene targets to inhibit HIV-1 viral reactivation. Separate but parallel approaches have been used in cancer contexts, whereby synthetic lethality is exploited to promote death of cancer cells. A recent study has led to identification of paralogs with redundant function that lead to cell death when a pair of gene paralogs are knocked out (58). From this study, 12% of paralogs tested lead to cell death in their context. We interpret this to mean that there is a great deal of gene paralogs which may serve redundant functions. Ongoing work will seek to identify factors that are like *CCNT1* in that when targeted, have drastic effects on viral replication, and minimal effects on the host – by focusing on top hits that have gene paralogs and thus may have redundancy. Other screen hits had Gene Effect scores similar to *CCNT1* – including *TBL1XR1*, *OTUD5*, and *AMBRA1* on the DepMap Portal (45), suggesting that these may either have paralogs or dispensable functions for cell biology.

While LPAs have been developed in a block and lock approach, this approach still remains a challenge. In the case of dCA – for instance –HIV confers resistance to this drug through mutations in the LTR, Nef and Vpr (59, 60). Targeting CCNT1 – or additional gene paralogs with redundant functions – may prove to be a strong compliment to these LPAs, given how drastic an affect *CCNT1* Knockouts have on HIV-1 replication. Although the shock and kill approach and discovery of LRAs has been a large area of focus in recent years, there may be a role for both approaches in permanently silencing the latent reservoirs in those tissue reservoirs which are resistant to LRAs. Further investigation of *CCNT1* knockout in macrophages, microglial cells and other resident tissues, as well as other genes which have redundancy in a similar regard as *CCNT1*, will provide a good path forward to identify additional block and lock mechanisms that may supplement other approaches to an HIV functional cure.

## Methods

### Cell Culture and Maintenance

HEK293T cells were cultured in DMEM (ThermoFisher, 11965092) along with Penicillin/Streptomycin (Pen/Strep) and 10% Fetal Bovine Serum (FBS). J-Lat cells were cultured in RPMI 1640 media (ThermoFisher, 11875093) supplemented with Pen/Strep, 10% Fetal Bovine Serum (FBS), and 10 mM HEPES (ThermoFisher, 15630080). Cells were maintained at 37°C with 5% CO_2_. Cells were routinely tested and found to be free of mycoplasma contamination. Primary CD4+ T cell media used was RPMI 1640 + 1x Anti-Anti (Gibco, 15240096), 1x GlutaMAX (ThermoFisher Scientific; 35050061), 10 mM HEPES, and 10% FBS.

#### HIV-CRISPR Library Transduction and Virus-Encapsidated CRISPR Guide Screening

The HIV-Dep library containing 525 genes (4191 sgRNAs) was previously described (17). For transduction of J-Lat cells, HEK293T cells were seeded in 20×6 well cell culture plates, transfected with the HIV-DEP plasmid (667 ng), psPax2 (GagPol, 500 ng), and MD2.G (VSVG, 333 ng) per well in 200 uL of serum-free DMEM (Thermo Fisher Scientific) along with 4.5 uL of TransIT-LT1 reagent (Mirus Bio LLC; MIR2305). VSVG pseudotyped lentivirus was harvested and filtered through a 0.22 um filter (Sigma-Aldrich, SE1M179M6). Virus was titered using TZM-bl (NIH AIDS Reagent Program; ARP-8129) cells. J-Lat 10.6 and J-Lat 5A8 previously knocked out for *ZAP* (12) were transduced with HIV-CRISPR library lentivirus with DEAE-Dextran (final concentration 20 ug/mL, Sigma-Aldrich; D9885) at a multiplicity of infection (MOI) of 0.5. After 24 hours, puromycin (Sigma, P8833) at a final concentration of 0.4 ug/mL was added to the culture to select for cells that received the vector. The screen was performed 11 days after transduction, by treating the HIV-Dep library transduced J-Lat cells with latency reversal agents AZD5582 1 nM (MedChemExpress, HY-12600) and I-BET151 2.5 uM (SelleckChem, S2780) or DMSO (Sigma, 472301) control. After 24 hours (day 12), the supernatants were harvested, filtered (Millipore Sigma, SE1M179M6), and loaded over a 20% sterile sucrose solution (20% sucrose, 1 mM EDTA, 20 mM HEPES, 100 mM NaCl, distilled water) placed on a prechilled SW32Ti rotor. The viral pellets were then concentrated at *70,000 x g* for 1 hour at 4°C and gently resuspended in 140 ul of DPBS (Gibco; 14190144) and allowed to resuspend overnight at 4°C. Simultaneously, transduced cells were harvested to isolate genomic DNA (gDNA). Cells were centrifuged and resuspended in DPBS. Cells were then spun down, supernatant removed, and cell pellets were frozen until ready for gDNA extraction.

### Latency HIV-CRISPR Screen

Viral RNA (vRNA) and gDNA was isolated as previously described (20). Briefly, vRNA was isolated using the QIAamp Viral RNA Mini Kit (Qiagen, 52904). Reverse transcription of vRNA was performed using SuperScript Reverse Transcriptase Kit (ThermoFisher, 18064014). gDNA was isolated using the QIAamp DNA Blood Midi Kit (Qiagen, 51183). vRNA and gDNA were both amplified by PCR using R1_forward primer and R1_Reverse primer using Herculase II Fusion DNA Polymerase (Agilent, 600677). PCR products were cleaned up using the QIAquick PCR clean up kit (Qiagen, 28104) and a second round of PCR was performed using R2_reverse primer and R2_IndexX primer (see supplementary file). The 230bp band was verified to be present and the amplified PCR products were cleaned up using double-sided SPRI via AMPure Beads (Beckman Coulter, A63880). Purified samples were normalized to a concentration of 10 nM using Qubit dsDNA HS Assay Kit (Invitrogen, Q32854) before sequencing.

Adapter sequences were computationally trimmed from sequencing results and the viral sequencing was compared relative to genomic knockout pool to determine the relative enrichment or depletion of each guide. An artificial NTC sgRNA gene set was generated that is equivalent to the number of genes present in the HIV-Dep library “synNTCs” by iteratively binning the NTC sgRNA sequences. MAGEcK and MAGEcK Flute statistical (22, 61) analyses were used to analyze the depletion of guides/genes in the RNA viral supernatent relative to their abundance in the cell DNA. Z-scores were determined as previously described (17, 62). For each HIV-Dep LAI replicate, and for each replicate of J-Lat CRISPR screen, z-scores were calculated. An average of the z-scores from each replicate was used to generate a heatmap. Heatmaps were generated using Morpheus (https://software.broadinstitute.org/morpheus). Code for z-score analysis of CRISPR screen data can be found at https://github.com/amcolash/hiv-crispr-zscore-analysis.

### Validation of Screen Hits

Genes identified in the HIV-Latency screen that were depleted after LRA treatment were validated either by lentiviral knockout or by electroporation of RNA guides and Cas9. For genes validated by lentiviral knockout, a forward and reverse primer corresponding with 2 individual guides targeting each gene were cloned into pLCV2 (see supplemental file for oligos used for each gene) and cells were transduced as described above. Puromycin selection continued for 10-14 days until treated with LRAs. For pooled electroporation knockout experiments, CRISPR/Cas9-mediated knockout was performed against genes of interest using Gene Knockout Kit v2 (Synthego). Guides targeting genes of interest (see supplemental file for guides used) with 1 uL of 20 uM Cas9-NLS (UC Berkeley Macro Lab) and RNP complexes were made with SE Cell Line 96-well Nucleofector Kit (Lonza, V4SC-1096). Complexes were incubated at room temperature for ten minutes, and 2E5 cells of J-Lat 10.6 were centrifuged at *100 x g* for 10 minutes at 25°C, and were resuspended in Cas9-RNP complexes and electroporated on Lonza 4D-Nucleofector using code CL-120. Cells were recovered with RPMI media pre-warmed to 37°C. Knockout pools were maintained for 10-14 days to allow for expansion and subsequently treated with LRAs. In both cases, reactivation was measured by RT activity as described (63) 24 hours after LRA treatment and genomic DNA analyzed to assess the degree of gene knockouts.

For *CCNT1* knockout clones, CRISPR/Cas9-mediated knockout was performed using Gene Knockout Kit v2 (Synthego). Guides targeting *CCNT1* were complexed with 1 uL of 20 uM Cas9-NLS (UC Berkeley Macro Lab) and RNP complexes were made with SE Cell Line 96-well Nucleofector Kit (Lonza, V4SC-1096). Complexes were incubated at room temperature for ten minutes, and 2E5 cells of J-Lat 10.6 were centrifuged at *100 x g* for 10 minutes at 25°C, and were resuspended in Cas9-RNP complexes and electroporated on Lonza 4D-Nucleofector using code CL-120. Cells were recovered with media pre-warmed to 37°C. Five days post-electroporation, single cells were sorted into a 96-well U-bottom plate filled with 100 uL RPMI media (20% FBS).

To assess the growth of *CCNT1* knockout J-Lat 10.6 relative to wild-type, three individual flasks of either wild-type, *CCNT1* Knockout 1 or *CCNT1* Knockout 2 J-Lat 10.6 were maintained for each line. Cells were resuspended at a concentration of 2E5 cells/mL in a total of 10 mL RPMI media. Cells were monitored and split approximately every two days. Cell counts prior to splitting were taken, the volume of cell suspension removed (the same volume was removed for each line) was tracked, and subtracted from overall cell count. These values were tracked over a span of nine days.

### Protein Isolation and Western Blotting

Cell pellets (1.5E6-3E6 cells) from pooled lentiviral knockout experiments (NTC10 and *CCNT1* sg1 and sg2) and clonal knockout experiments (J-Lat 10.6 *CCNT1* KO clone 1 and 2) were isolated from each respective experiment. Supernatant was removed and cells were resuspended in 500 uL of cold (4°C) 1x PBS. Cells were pelleted, resuspended in 100 uL of RIPA buffer (150 mM NaCl (Sigma, S3014), 50mM Tris pH 8.0, 1% NP-40 (Calbiochem, 492016), 0.5% Sodium Deoxycholate (Sigma-Aldrich, D6750), and 0.1% SDS (Sigma-Aldrich, L4509), Benzonase 1 uL/mL (Millipore, 70664), and cOmplete Protease Inhibitor Cocktail (Roche; 11697498001), and incubated on ice for 10 minutes with repeated vortexing. Cell lysate was pelleted at *20,000 x g* for 20 minutes at 4°C. Clarified supernatant was transferred to a new tube and quantified by BCA. Samples were prepared by adding 4x NuPAGE LDS Sample Buffer (ThermoFisher, NP0007) with 5% 2-Mercaptoethanol (Sigma-Aldrich, M3148) and denatured at 95°C for 5 minutes. Lysates were run on a NuPAGE 4-12% Bis-Tris pre-cast gel (ThermoFisher Scientific; NP0336) and transferred to a nitrocellulose membrane (Biorad; 1620115). After transfer, nitrocellulose membrane was blocked in 0.1% Tween/5% Milk in 1XPBS solution for 30 minutes at room temperature. Primary antibodies used for western blotting were mouse α-CCNT1 (Santa Cruz Biotechnology, sc-271348, 1:500), mouse α-CCNT2 (Santa Cruz Biotechnology, sc-81243, 1:500), and rabbit α-actin (Sigma-Aldrich, A2066 1:5000). Antibodies were diluted in 1x PBS-Tween 0.1% (PBST) and rocked on nitrocellulose membrane overnight at 4°C. Membrane was washed with PBST 3-5 times, for 5 minutes each wash. The following secondary antibody dilutions were made 1:2000 in PBST: goat α-mouse IgG-HRP (R&D Systems; HAF007) and goat α-rabbit IgG-HRP (R&D Systems; HAF008). SuperSignal West Femto Maximum Sensitivity Substrate (ThermoFisher; 34095) was used for CCNT1 and CCNT2, and SuperSignal West Pico PLUS Chemiluminescent Substrate (ThermoFisher, 34580) was used for Actin. Visualization was done on a BioRad Chemidoc MP Imaging System.

### Genomic Editing Analysis

Cells for each knockout were pelleted, washed with 1X PBS, supernatant removed, and cell pellets frozen at -80°C until ready for DNA isolation. Genomic DNA was isolated using QIAamp DNA Blood Mini Kit (Qiagen; 51104). The gene of interest was amplified using primers described using either Q5 High-Fidelity DNA polymerase (NEB; M0491S) or Platinum Taq DNA polymerase High Fidelity (ThermoFisher Scientific; 11304011). PCR products were purified using AMPure beads (Beckman Coulter, A63880) or QIAquick PCR clean up kit (Qiagen, 28104) and submitted to Fred Hutch Genomics shared resource for sequencing. Analysis was performed using Inference of CRISPR Edits (ICE) (64).

### LRA Treatments

For J-Lat 10.6 or J-Lat 5A8 cells, LRAs were used at the following concentrations: TNFα (Peprotech, 300-01A) 10 ng/mL; AZD5582 (MedChemExpress, HY-12600) 1 nM; I-BET151 (SelleckChem, S2780) 2.5 uM; Prostratin (Sigma-Aldrich, P0077) 0.1 uM; SAHA/Vorinostat (SelleckChem, S1047), 2.5 uM. For CD3/CD28 antibody stimulation Anti-CD3 clone UCHT1 (Tonbo, 40-0038-U500) was plated on 96-well flat bottom plate at 10 ug/mL in 1x PBS, incubated overnight at 4°C, aspirated, and CD28 clone 28.2 antibody (Tonbo, 40-0289-U500) was added to RPMI media at a concentration of 4 ug/mL for cell resuspension. Cells for each experiment were resuspended at a concentration of 5E5 cells/mL in appropriate LRA media, and 200 uL was aliquoted into 96-well flat blottom TC plate. For Primary CD4+ T Cell LRA treatment, PMA (Sigma-Aldrich, P1585) was used at a concentration of 10 nM, in combination with ionomycin (Sigma-Aldrich, I0634) was used at a concentration of 1 uM. For primary cell experiments, CD3 antibody (Tonbo, 40-0038-U500) was used at a concentration of 10 ug/mL and CD28 antibody (Tonbo, 40-0289-U500) at a concentration of 5 ug/mL. All LRA treatments were performed for 24 hours unless otherwise indicated.

### Primary CD4+ Cell Isolation and Latency Model

All centrifugation steps of Primary CD4+ T cells were performed at *300 x g* for 10 minutes at 25°C unless otherwise noted. PBMCs were isolated from used leukocyte filters (Bloodworks Northwest) over a Ficoll gradient (Millipore Sigma, GE17-1440-02), cryofrozen at a concentration of 10-20E6 cells/mL in 90% FBS/10% DMSO, and stored in liquid nitrogen until ready to use. On thawing, PBMCs were washed dropwise with pre-warmed RPMI-1640 media (Thermo Fisher) and treated with benzonase (25 U/mL) (Sigma-Aldrich, E1014) for 15 minutes at room temperature. PBMCs were maintained at a concentration of 2E6 cells/mL overnight at 37°C. The following day, CD4+ T cells were isolated using the EasySep Human CD4+ T cell Isolation Kit (Stemcell Technologies, 17952) and subsequently activated using the T Cell Activation/Expansion Kit (Miltenyi Biotec, 130-091-441). From this point forward, CD4+ T cells were cultured in RPMI + IL-2 (final concentration 100 U/mL, Roche, 10799068001), IL-7 (final conc. 2 ng/mL, Peprotech, 200-07) and IL-15 (final conc. 2 ng/mL, Peprotech, 200-15) unless otherwise noted. Cells were activated continually for two days prior to infection.

Lentivirus for infection of primary CD4+ T cells was generated by transfecting HEK293T cells with *Δ6-dGFP-Thy1.2-Gagpol+* Plasmid (900 ng, gift from Ed Browne Lab), psPax2 plasmid (450 ng), and MD2.Cocal plasmid (150 ng, gift from Hans-Peter Kiem Lab (65). After two days, virus was filtered using a millipore filter (Millipore Sigma, SE1M179M6). On day of infection, activation beads were first magnetically removed. Infection of CD4+ T cells was performed by aliquoting 5E6 CD4+ T cells iteratively into 50 mL falcon tubes, and resuspending in virus + polybrene (final conc 8 ug/mL, Sigma-Aldrich, TR-1003) or RPMI media + polybrene for the uninfected control at a concentration of 1E6 cells/mL. Spinoculation was performed for *1100 x g* for 2 hours at 30°C. Cells were maintained at a concentration of 1E6 cells/mL.

Three days post-infection, a small portion of cells were taken to assess infection by staining with CD90-AF700 antibody (Biolegend, 140323) for 20 minutes (1:1000 dilution in FACS Buffer), fixing with 4% paraformaldehyde and sorting by AF700 and GFP on SP Celesta 2 Cell Analysis Machine (Flow Cytometry Core, Fred Hutch). CD90+ cells were then isolated using the CD90.2 Cell Isolation Kit (Stemcell Technologies, 18951). Two days after CD90+ cells were purified, cells then were electroporated using electroporation code EH-100 and using the P3 Primary Cell 96-well Nucleofector Kit (Lonza, V4SP-3096). Knockout pools were maintained for an additional nine days prior to coculturing with H80 feeder cell line with IL-2 (Final conc 20 U/mL) in RPMI (no longer cultured with IL-7 and IL-15). Four days later, the cells were treated with PMAi or CD3/CD28 antibody co-stimulation (or unstimulated control) and analyzed on SP Celesta 2 (Core Facility) to evaluate reactivation potential by assessing Thy1.2, CD90+ and GFP+ cells. An early activation marker of T cells was also monitored using PE-Conjugated CD-69 antibody (Biolegend, 310906). Analysis was performed on FlowJo software. Genomic DNA was isolated at the end of experiment from uninfected and knockout cells to assess for genomic ICE analysis.

### Primary CD4+ T cell activation Test

CD4+ cells were isolated from healthy donors and activated as described above. After two days of activation, beads were magnetically removed. Three days later, cells electroporated following the protocol above, and treated with CD3/CD28 antibody after cells were allowed to recover for two additional days. Activation was monitored using PE-Conjugated CD69 antibody (Biolegend, 310906) on SP Celesta 2. Genomic DNA was isolated for analysis.

### RNA-seq analysis of *CCNT1* knockout cells

For RNA isolated from J-Lat 10.6 cells, cells first were passaged and split equally three times prior to isolation. J-Lat 10.6 either wild-type for *CCNT1* or knocked out for *CCNT1* were each treated with TNFα (Peprotech, 300-01A) at 10 ng/mL or unstimulated in triplicate. For primary cell experiments, knockouts were performed similarly as described in “Primary CD4+ T cell activation Test,” and RNA was isolated after LRA treatment. In both J-Lat and primary CD4+ T cell isolation experiments, 0.1-2E6 cells were isolated and resuspended in 350 uL of RLT Plus (Qiagen, 1053393) + 1% 2-mercaptoethanol (Millipore Sigma, M3148). Cells were frozen in buffer RLT plus until ready to continue with isolation. Thawed RLT lysates were then run over a QIAshredder column (Qiagen, 79654) and subsequently over a gDNA eliminator column. Qiagen RNeasy Plus Mini Kit was then used in order to obtain purified total RNA. RNA was submitted for TapeStation RNA assay or HighSense RNA assay (Fred Hutch Core Facilities) and RINe scores were all found to be ≥ 9.6.

### RNAseq Analysis Methods

Quality assessment of the raw sequencing data, in Fastq format, was performed with fastp v0.20.0 (66) to ensure that data had high base call quality, expected GC content for RNA-seq, and no overrepresented contaminating sequences. No reads or individual bases were removed during this assessment step. The fastq files were aligned to the UCSC human hg38 reference assembly using STAR v2.7.7 (67). STAR was run with the parameter “—quantMode GeneCounts” to produce a table of raw gene-level counts with respect to annotations from human GENCODE build v38. To account for unstranded library preparation, only unstranded counts from the table were retained for further analysis. The quality of the alignments was evaluated using RSeQC v3.0.0 (68) including assessment of bam statistics, read-pair inner distance, and read distribution. Differential expression analysis was performed with edgeR v3.36.0 (69) to identify the differences between knockout stimulated and stimulated for with *CCNT1* and *AAVS1* genes, as well as differences between the two genes in knockout and knockout stimulated conditions. Genes with very low expression across all samples were flagged for removal by filterbyExpr, and TMM normalization was applied with calcNormFactors to account for differences in library composition and sequencing depth. We constructed a design matrix to incorporate potential batch effects related to donor information, after which the dispersion of expression values was estimated using estimateDisp. Testing for each gene was then performed with the QL F-test framework using glmQLFTest which outputs for each gene a p-value, a log2(fold change) value, and a Benjamini-Hochberg corrected false discovery rate (FDR) to control for multiple-testing. The results were plotted using ggplot2 v3.3.5 (70). For analysis of J-Lat 10.6 RNA sequencing data, we used the reference genome previously assembled and described for J-Lat 10.6 (12). Using this reference, we masked the 5’ LTR of the integrated provirus. All splice variants as well as genomic RNA that terminate at a polyA site in the 3’ LTR are similarly named “HIV-1.”

## Acknowledgments

We thank Molly OhAinle, Joy Twentyman, and Carley Gray for critical feedback of this manuscript, and members of the Emerman lab for helpful suggestions and technical assistance. We thank Warner Greene at Gladstone Institute of Virology and Immunology and University of California, San Francisco, San Francisco, CA, USA for sharing the J-Lat 5A8 cells and Darell Bigner at Duke University, Durham, NC, USA for sharing the H80 cells. Harini Srinivasan and Matt Fitzgibbon at the FHCC for bioinformatics support. This work was supported by DP1 DA051110 (ME), and CARE 1UM1-A1-164567 (ME). Y.D. was supported by NIH grant R25 GM086304. This research was also supported by the Genomics, Bioinformatics, and Flow Cytometry Shared Resource, RRID:SCR_022606, of the Fred Hutch/University of Washington Cancer Consortium (P30 CA015704).

## Supplemental figure legends

**Supplemental Figure S1: Supplement to figure 3.** Flow plots are shown from one representative donor. CD4+ T Cells infected with dual-reporter HIV virus, gene knockouts performed, latency was established, and treated with anti-CD3/anti-CD28 antibodies for 24 hours (see Methods) to assess reactivation potential with T cell receptor co-stimulation.

## Supplemental Files Description

**Supplemental File 1:** The latency HIV-CRISPR screen results of the J-Lat 10.6 (Sheet 1) and J-Lat 5A8 line (Sheet 2) are shown in ranked order of the most depleted guides. The mean z-scores of the Jurkat LAI screen (previous study, 4 replicates) and J-Lat 10.6 and 5A8 screens (2 replicates) are shown. Related to Figure 1 and Figure 2.

**Supplemental File 2:** ICE genomic analysis of *AAVS1* and *CCNT1* knockouts is shown for each donor of the primary CD4+ T cell RNA sequencing experiment. Related to Figure 5.

## Notes

### Competing Interest Statement

The authors have declared no competing interest.

## References

1. Kim Y, Anderson JL, Lewin SR. 2018. Getting the “Kill” into “Shock and Kill”: Strategies to Eliminate Latent HIV. Cell Host Microbe 23:14–26.

2. Rodari A, Darcis G, Van Lint CM. 2021. The Current Status of Latency Reversing Agents for HIV-1 Remission. Annu Rev Virol 8:491–514.

3. Margolis DM, Archin NM, Cohen MS, Eron JJ, Ferrari G, Garcia JV, Gay CL, Goonetilleke N, Joseph SB, Swanstrom R, Turner AW, Wahl A. 2020. Curing HIV: Seeking to Target and Clear Persistent Infection. Cell 181:189–206.

4. Cohn LB, Chomont N, Deeks SG. 2020. The Biology of the HIV-1 Latent Reservoir and Implications for Cure Strategies. Cell Host Microbe 27:519–530.

5. Grau-Exposito J, Luque-Ballesteros L, Navarro J, Curran A, Burgos J, Ribera E, Torrella A, Planas B, Badia R, Martin-Castillo M, Fernandez-Sojo J, Genesca M, Falco V, Buzon MJ. 2019. Latency reversal agents affect differently the latent reservoir present in distinct CD4+ T subpopulations. PLoS Pathog 15:e1007991.

6. Li C, Mori L, Valente ST. 2021. The Block-and-Lock Strategy for Human Immunodeficiency Virus Cure: Lessons Learned from Didehydro-Cortistatin A. J Infect Dis 223:46–53.

7. Vansant G, Bruggemans A, Janssens J, Debyser Z. 2020. Block-And-Lock Strategies to Cure HIV Infection. Viruses 12.

8. Mediouni S, Chinthalapudi K, Ekka MK, Usui I, Jablonski JA, Clementz MA, Mousseau G, Nowak J, Macherla VR, Beverage JN, Esquenazi E, Baran P, de Vera IMS, Kojetin D, Loret EP, Nettles K, Maiti S, Izard T, Valente ST. 2019. Didehydro-Cortistatin A Inhibits HIV-1 by Specifically Binding to the Unstructured Basic Region of Tat. mBio 10.

9. Turner AM, Ackley AM, Matrone MA, Morris KV. 2012. Characterization of an HIV-targeted transcriptional gene-silencing RNA in primary cells. Hum Gene Ther 23:473–83.

10. Ahlenstiel C, Mendez C, Lim ST, Marks K, Turville S, Cooper DA, Kelleher AD, Suzuki K. 2015. Novel RNA Duplex Locks HIV-1 in a Latent State via Chromatin-mediated Transcriptional Silencing. Mol Ther Nucleic Acids 4:e261.

11. Marconi VC, Moser C, Gavegnano C, Deeks SG, Lederman MM, Overton ET, Tsibris A, Hunt PW, Kantor A, Sekaly RP, Tressler R, Flexner C, Hurwitz SJ, Moisi D, Clagett B, Hardin WR, Del Rio C, Schinazi RF, Lennox JJ. 2022. Randomized Trial of Ruxolitinib in Antiretroviral-Treated Adults With Human Immunodeficiency Virus. Clin Infect Dis 74:95–104.

12. Hsieh E, Janssens DH, Paddison PJ, Browne EP, Henikoff S, OhAinle M, Emerman M. 2023. A modular CRISPR screen identifies individual and combination pathways contributing to HIV-1 latency. PLoS Pathog 19:e1011101.

13. Bacon CW, D’Orso I. 2019. CDK9: a signaling hub for transcriptional control. Transcription 10:57–75.

14. Peng J, Zhu Y, Milton JT, Price DH. 1998. Identification of multiple cyclin subunits of human P-TEFb. Genes Dev 12:755–62.

15. Lin X, Taube R, Fujinaga K, Peterlin BM. 2002. P-TEFb containing cyclin K and Cdk9 can activate transcription via RNA. J Biol Chem 277:16873–8.

16. Bieniasz PD,, Grdina TA, Bogerd HP, Cullen BR,. 1999. Analysis of the Effect of Natural Sequence Variation in Tat and in Cyclin T on the Formation and RNA Binding Properties of Tat-Cyclin T Complexes. 73:5777–5786.

17. Montoya VR, Ready TM, Felton A, Fine SR, OhAinle M, Emerman M. 2023. A Virus-Packageable CRISPR System Identifies Host Dependency Factors Co-Opted by Multiple HIV-1 Strains. mBio 14:e0000923.

18. Zhou Y, Zhou B, Pache L, Chang M, Khodabakhshi AH, Tanaseichuk O, Benner C, Chanda SK. 2019. Metascape provides a biologist-oriented resource for the analysis of systems-level datasets. Nat Commun 10:1523.

19. OhAinle M, Helms L, Vermeire J, Roesch F, Humes D, Basom R, Delrow JJ, Overbaugh J, Emerman M. 2018. A virus-packageable CRISPR screen identifies host factors mediating interferon inhibition of HIV. Elife 7.

20. Roesch F, OhAinle M. 2020. HIV-CRISPR: A CRISPR/Cas9 Screening Method to Identify Genes Affecting HIV Replication. Bio Protoc 10:e3614.

21. Falcinelli SD, Peterson JJ, Turner AW, Irlbeck D, Read J, Raines SL, James KS, Sutton C, Sanchez A, Emery A, Sampey G, Ferris R, Allard B, Ghofrani S, Kirchherr JL, Baker C, Kuruc JD, Gay CL, James LI, Wu G, Zuck P, Rioja I, Furze RC, Prinjha RK, Howell BJ, Swanstrom R, Browne EP, Strahl BD, Dunham RM, Archin NM, Margolis DM. 2022. Combined noncanonical NF-kappaB agonism and targeted BET bromodomain inhibition reverse HIV latency ex vivo. J Clin Invest 132.

22. Wang B, Wang M, Zhang W, Xiao T, Chen CH, Wu A, Wu F, Traugh N, Wang X, Li Z, Mei S, Cui Y, Shi S, Lipp JJ, Hinterndorfer M, Zuber J, Brown M, Li W, Liu XS. 2019. Integrative analysis of pooled CRISPR genetic screens using MAGeCKFlute. Nat Protoc 14:756–780.

23. Chu VT, Weber T, Wefers B, Wurst W, Sander S, Rajewsky K, Kuhn R. 2015. Increasing the efficiency of homology-directed repair for CRISPR-Cas9-induced precise gene editing in mammalian cells. Nat Biotechnol 33:543–8.

24. Ivanov D, Kwak YT, Nee E, Guo J, Garcia-Martinez LF, Gaynor RB. 1999. Cyclin T1 domains involved in complex formation with Tat and TAR RNA are critical for tat-activation. J Mol Biol 288:41–56.

25. Jones KA. 1997. Taking a new TAK on tat transactivation. Genes Dev 11:2593–9.

26. Jones KA, Peterlin BM. 1994. Control of RNA initiation and elongation at the HIV-1 promoter. Annu Rev Biochem 63:717–43.

27. Kazanietz MG, Areces LB, Bahador A, Mischak H, Goodnight J, Mushinski JF, Blumberg PM. 1993. Characterization of ligand and substrate specificity for the calcium-dependent and calcium-independent protein kinase C isozymes. Mol Pharmacol 44:298–307.

28. Sung TL, Rice AP. 2006. Effects of prostratin on Cyclin T1/P-TEFb function and the gene expression profile in primary resting CD4+ T cells. Retrovirology 3:66.

29. Elliott JH, Wightman F, Solomon A, Ghneim K, Ahlers J, Cameron MJ, Smith MZ, Spelman T, McMahon J, Velayudham P, Brown G, Roney J, Watson J, Prince MH, Hoy JF, Chomont N, Fromentin R, Procopio FA, Zeidan J, Palmer S, Odevall L, Johnstone RW, Martin BP, Sinclair E, Deeks SG, Hazuda DJ, Cameron PU, Sekaly RP, Lewin SR. 2014. Activation of HIV transcription with short-course vorinostat in HIV-infected patients on suppressive antiretroviral therapy. PLoS Pathog 10:e1004473.

30. Fujinaga K, Huang F, Peterlin BM. 2023. P-TEFb: The master regulator of transcription elongation. Mol Cell 83:393–403.

31. Bauer D, Mazzio E, Hilliard A, Oriaku ET, Soliman KFA. 2020. Effect of apigenin on whole transcriptome profile of TNFalpha-activated MDA-MB-468 triple negative breast cancer cells. Oncol Lett 19:2123–2132.

32. Oeckinghaus A, Ghosh S. 2009. The NF-kappaB family of transcription factors and its regulation. Cold Spring Harb Perspect Biol 1:a000034.

33. Palmer S, Chen YH. 2008. Bcl-3, a multifaceted modulator of NF-kappaB-mediated gene transcription. Immunol Res 42:210–8.

34. Dillon SR, Sprecher C, Hammond A, Bilsborough J, Rosenfeld-Franklin M, Presnell SR, Haugen HS, Maurer M, Harder B, Johnston J, Bort S, Mudri S, Kuijper JL, Bukowski T, Shea P, Dong DL, Dasovich M, Grant FJ, Lockwood L, Levin SD, LeCiel C, Waggie K, Day H, Topouzis S, Kramer J, Kuestner R, Chen Z, Foster D, Parrish-Novak J, Gross JA. 2004. Interleukin 31, a cytokine produced by activated T cells, induces dermatitis in mice. Nat Immunol 5:752–60.

35. Kohoutek J, Li Q, Blazek D, Luo Z, Jiang H, Peterlin BM. 2009. Cyclin T2 is essential for mouse embryogenesis. Mol Cell Biol 29:3280–5.

36. Oven I, Brdickova N, Kohoutek J, Vaupotic T, Narat M, Peterlin BM. 2007. AIRE recruits P-TEFb for transcriptional elongation of target genes in medullary thymic epithelial cells. Mol Cell Biol 27:8815–23.

37. Ramakrishnan R, Yu W, Rice AP,. 2011. Limited redundancy in genes regulated by Cyclin T2 and Cyclin T1. 4.

38. Heisterkamp N, Groffen J, Warburton D, Sneddon TP. 2008. The human gamma-glutamyltransferase gene family. Hum Genet 123:321–32.

39. Berg JS, Derfler BH, Pennisi CM, Corey DP, Cheney RE. 2000. Myosin-X, a novel myosin with pleckstrin homology domains, associates with regions of dynamic actin. J Cell Sci 113 Pt 19:3439–51.

40. Uhl J, Gujarathi S, Waheed AA, Gordon A, Freed EO, Gousset K. 2019. Myosin-X is essential to the intercellular spread of HIV-1 Nef through tunneling nanotubes. J Cell Commun Signal 13:209–224.

41. Zhang S, Laouar A, Denzin LK, Sant’Angelo DB. 2015. Zbtb16 (PLZF) is stably suppressed and not inducible in non-innate T cells via T cell receptor-mediated signaling. Sci Rep 5:12113.

42. Yasuda-Yamahara M, Rogg M, Yamahara K, Maier JI, Huber TB, Schell C. 2018. AIF1L regulates actomyosin contractility and filopodial extensions in human podocytes. PLoS One 13:e0200487.

43. Davis DB, Delmonte AJ, Ly CT, McNally EM. 2000. Myoferlin, a candidate gene and potential modifier of muscular dystrophy. Hum Mol Genet 9:217–26.

44. Bernatchez PN, Acevedo L, Fernandez-Hernando C, Murata T, Chalouni C, Kim J, Erdjument-Bromage H, Shah V, Gratton JP, McNally EM, Tempst P, Sessa WC. 2007. Myoferlin regulates vascular endothelial growth factor receptor-2 stability and function. J Biol Chem 282:30745–53.

45. Tsherniak A, Vazquez F, Montgomery PG, Weir BA, Kryukov G, Cowley GS, Gill S, Harrington WF, Pantel S, Krill-Burger JM, Meyers RM, Ali L, Goodale A, Lee Y, Jiang G, Hsiao J, Gerath WFJ, Howell S, Merkel E, Ghandi M, Garraway LA, Root DE, Golub TR, Boehm JS, Hahn WC. 2017. Defining a Cancer Dependency Map. Cell 170:564–576 e16.

46. Ghose R, Liou LY, Herrmann CH, Rice AP. 2001. Induction of TAK (cyclin T1/P-TEFb) in purified resting CD4(+) T lymphocytes by combination of cytokines. J Virol 75:11336–43.

47. Garriga J, Peng J, Parreno M, Price DH, Henderson EE, Grana X. 1998. Upregulation of cyclin T1/CDK9 complexes during T cell activation. Oncogene 17:3093–102.

48. Huang F, Nguyen TT, Echeverria I, Rakesh R, Cary DC, Paculova H, Sali A, Weiss A, Peterlin BM, Fujinaga K. 2021. Reversible phosphorylation of cyclin T1 promotes assembly and stability of P-TEFb. Elife 10.

49. Guenther MG, Lane WS, Fischle W, Verdin E, Lazar MA, Shiekhattar R. 2000. A core SMRT corepressor complex containing HDAC3 and TBL1, a WD40-repeat protein linked to deafness. Genes Dev 14:1048–57.

50. Ning L, Rui X, Bo W, Qing G. 2021. The critical roles of histone deacetylase 3 in the pathogenesis of solid organ injury. Cell Death Dis 12:734.

51. Yoon HG, Chan DW, Huang ZQ, Li J, Fondell JD, Qin J, Wong J. 2003. Purification and functional characterization of the human N-CoR complex: the roles of HDAC3, TBL1 and TBLR1. EMBO J 22:1336-46.

52. Marks PA, Breslow R. 2007. Dimethyl sulfoxide to vorinostat: development of this histone deacetylase inhibitor as an anticancer drug. Nat Biotechnol 25:84–90.

53. Dai W, Wu F, McMyn N, Song B, Walker-Sperling VE, Varriale J, Zhang H, Barouch DH, Siliciano JD, Li W, Siliciano RF. 2022. Genome-wide CRISPR screens identify combinations of candidate latency reversing agents for targeting the latent HIV-1 reservoir. Sci Transl Med 14:eabh3351.

54. Zhou L, Jiang Y, Luo Q, Li L, Jia L. 2019. Neddylation: a novel modulator of the tumor microenvironment. Mol Cancer 18:77.

55. Liang WS, Maddukuri A, Teslovich TM, de la Fuente C, Agbottah E, Dadgar S, Kehn K, Hautaniemi S, Pumfery A, Stephan DA, Kashanchi F. 2005. Therapeutic targets for HIV-1 infection in the host proteome. Retrovirology 2:20.

56. Hyrcza MD, Kovacs C, Loutfy M, Halpenny R, Heisler L, Yang S, Wilkins O, Ostrowski M, Der SD. 2007. Distinct transcriptional profiles in ex vivo CD4+ and CD8+ T cells are established early in human immunodeficiency virus type 1 infection and are characterized by a chronic interferon response as well as extensive transcriptional changes in CD8+ T cells. J Virol 81:3477–86.

57. Dixit U, Bhutoria S, Wu X, Qiu L, Spira M, Mathew S, Harris R, Adams LJ, Cahill S, Pathak R, Rajesh Kumar P, Nguyen M, Acharya SA, Brenowitz M, Almo SC, Zou X, Steven AC, Cowburn D, Girvin M, Kalpana GV. 2021. INI1/SMARCB1 Rpt1 domain mimics TAR RNA in binding to integrase to facilitate HIV-1 replication. Nat Commun 12:2743.

58. Parrish PCR, Thomas JD, Gabel AM, Kamlapurkar S, Bradley RK, Berger AH. 2021. Discovery of synthetic lethal and tumor suppressor paralog pairs in the human genome. Cell Rep 36:109597.

59. Mousseau G, Aneja R, Clementz MA, Mediouni S, Lima NS, Haregot A, Kessing CF, Jablonski JA, Thenin-Houssier S, Nagarsheth N, Trautmann L, Valente ST. 2019. Resistance to the Tat Inhibitor Didehydro-Cortistatin A Is Mediated by Heightened Basal HIV-1 Transcription. mBio 10.

60. Rice AP. 2019. Unexpected Mutations in HIV-1 That Confer Resistance to the Tat Inhibitor Didehydro-Cortistatin A. mBio 10.

61. Li W, Xu H, Xiao T, Cong L, Love MI, Zhang F, Irizarry RA, Liu JS, Brown M, Liu XS. 2014. MAGeCK enables robust identification of essential genes from genome-scale CRISPR/Cas9 knockout screens. Genome Biol 15:554.

62. Schneider WM, Luna JM, Hoffmann HH, Sanchez-Rivera FJ, Leal AA, Ashbrook AW, Le Pen J, Ricardo-Lax I, Michailidis E, Peace A, Stenzel AF, Lowe SW, MacDonald MR, Rice CM, Poirier JT. 2021. Genome-Scale Identification of SARS-CoV-2 and Pan-coronavirus Host Factor Networks. Cell 184:120–132 e14.

63. Vermeire J, Naessens E, Vanderstraeten H, Landi A, Iannucci V, Van Nuffel A, Taghon T, Pizzato M, Verhasselt B. 2012. Quantification of reverse transcriptase activity by real-time PCR as a fast and accurate method for titration of HIV, lenti-and retroviral vectors. PLoS One 7:e50859.

64. Conant D, Hsiau T, Rossi N, Oki J, Maures T, Waite K, Yang J, Joshi S, Kelso R, Holden K, Enzmann BL, Stoner R. 2022. Inference of CRISPR Edits from Sanger Trace Data. CRISPR J 5:123–130.

65. Humbert O, Gisch DW, Wohlfahrt ME, Adams AB, Greenberg PD, Schmitt TM, Trobridge GD, Kiem HP. 2016. Development of Third-generation Cocal Envelope Producer Cell Lines for Robust Lentiviral Gene Transfer into Hematopoietic Stem Cells and T-cells. Mol Ther 24:1237–46.

66. Chen S, Zhou Y, Chen Y, Gu J. 2018. fastp: an ultra-fast all-in-one FASTQ preprocessor. Bioinformatics 34:i884–i890.

67. Dobin A, Davis CA, Schlesinger F, Drenkow J, Zaleski C, Jha S, Batut P, Chaisson M, Gingeras TR. 2013. STAR: ultrafast universal RNA-seq aligner. Bioinformatics 29:15–21.

68. Wang L, Wang S, Li W. 2012. RSeQC: quality control of RNA-seq experiments. Bioinformatics 28:2184–5.

69. Robinson MD, McCarthy DJ, Smyth GK. 2010. edgeR: a Bioconductor package for differential expression analysis of digital gene expression data. Bioinformatics 26:139–40.

70. Wickham H. 2016. ggplot2: Elegant Graphics for Data Analysis. Springer-Verlag New York.

