## Supplementary figures and images for "A CRISPR screen of HIV dependency factors reveals *CCNT1* is non-essential in T cells but required for HIV-1 reactivation from latency"

### SuppFig1_v2_072623.pdf

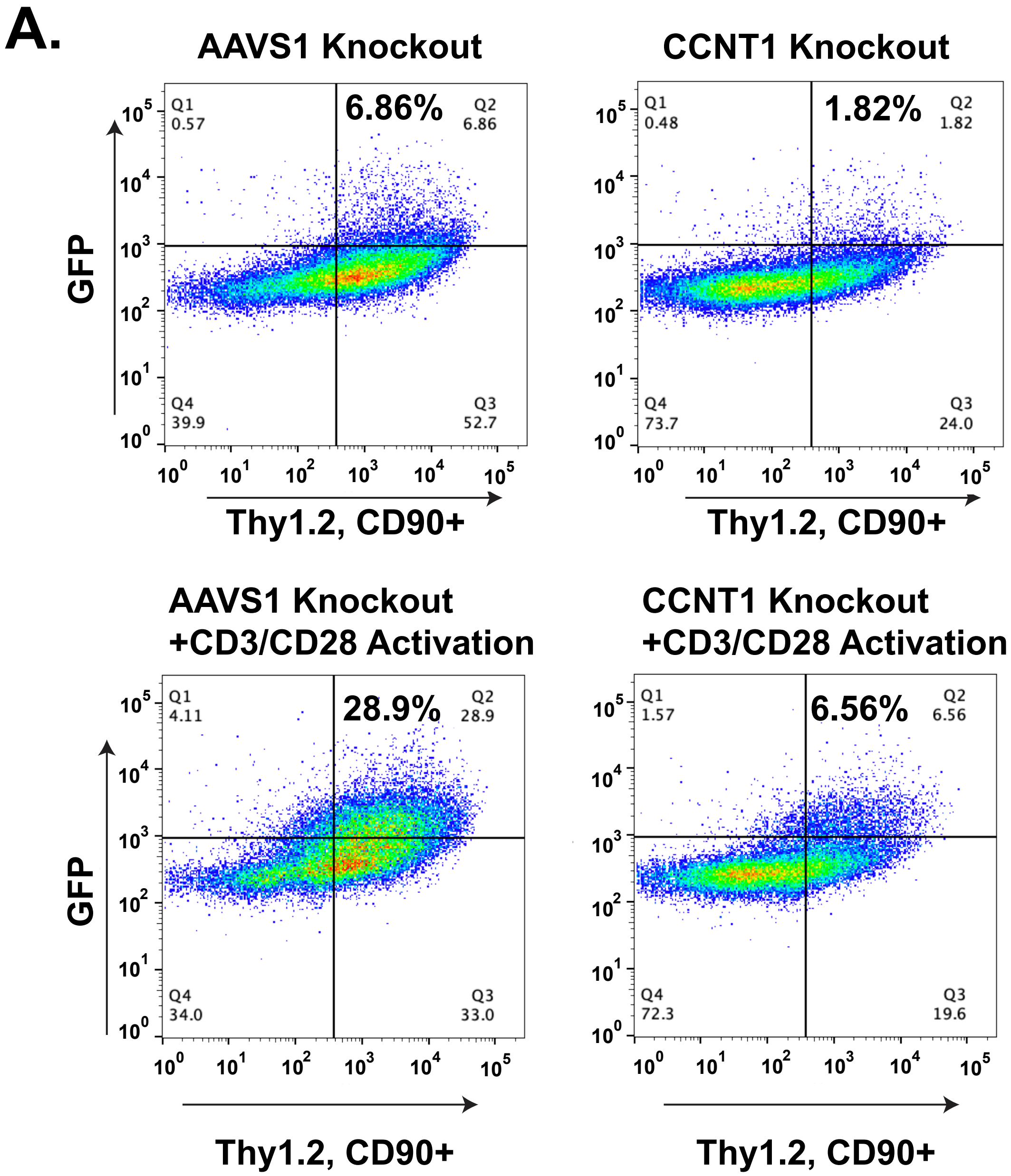
